# Substrate recognition, not sequestration, drives the engagement of an H3K9 methyltransferase in living cells

**DOI:** 10.64898/2026.06.30.735653

**Authors:** Yekaterina Fyodorova, Samuel B. Steen, Alexander Levashkevich, Luis A. Ortiz-Rodríguez, Sumanth Kumar Maheshwaram, Serena Chiu, Kaushik Ragunathan, Julie S. Biteen

## Abstract

The histone H3K9 methyltransferase Clr4 is essential for heterochromatin formation in *Schizosaccharomyces pombe*, yet how it searches for and engages chromatin *in vivo* remains unclear. Using live-cell single-molecule tracking of PAmCherry-Clr4, we quantified how Clr4 mutations alter its diffusion, search behavior, and residence times. Chromodomain and SET-domain Clr4 mutants move faster and more isotropically, reflecting impaired substrate recognition. In contrast, deleting Swi6, which has been proposed to sequester Clr4, does not reduce the slow-state fraction or alter Clr4 diffusion, indicating that chromatin engagement is intrinsic to Clr4 rather than HP1-dependent. Anisotropy analysis at short and intermediate displacements indicates that Clr4 does not explore chromatin by simple three-dimensional diffusion but through a guided, distance-dependent search in which it repeatedly samples nearby nucleosomes before disengaging. Across all other perturbations we examined, such as deletion of the CLRC component Rik1, impaired Clr4 ubiquitination, and using cells with an unmethylatable H3K9R substrate, Clr4 dynamics were only modestly affected, and a chromatin-associated population persisted in every background. This robustness indicates that the chromodomain and SET domains are the primary determinants of how Clr4 engages chromatin *in vivo*, allowing it to continuously sample the genome while maintaining a stable bound population. Our results suggest that the promiscuous sampling of chromatin may also enable Clr4 to establish heterochromatin *de novo* during adaptation.

**Table of Contents Graphic:** 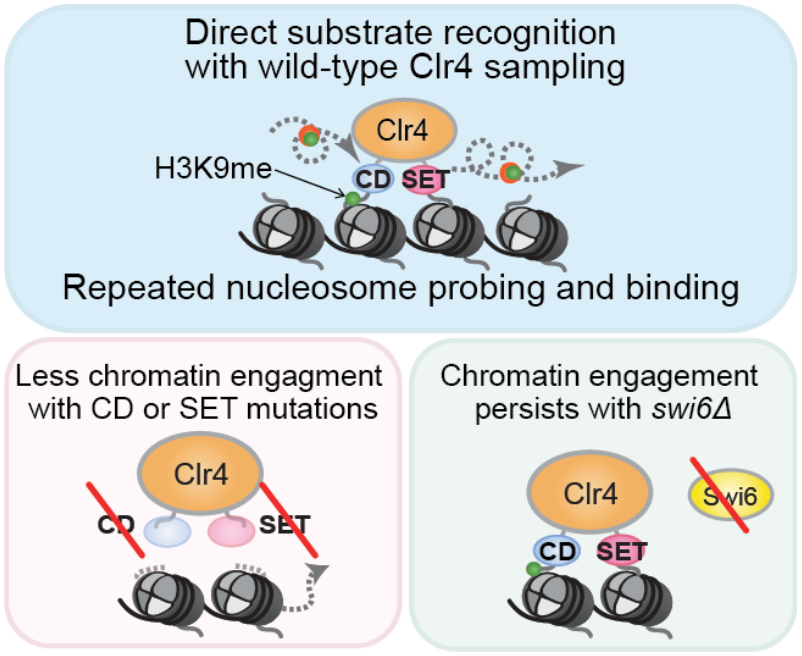

## Introduction

Epigenetic modifications regulate gene expression states without altering the primary DNA sequence. This process is vital in contexts such as multicellularity and cell fate specification, where cells with otherwise identical genomes must establish and maintain distinct phenotypic states.^1^ The covalent modification of DNA packaging proteins called histones contributes to this process. Histone methylation at specific lysine residues provides binding sites for chromatin-associated “reader” and “writer” proteins that reversibly affect transcription. Remarkably, the same chemical modification (lysine methylation) produces different transcriptional outcomes depending on its location: H3K4 methylation drives the recruitment of factors that promote transcription, whereas H3K9 methylation recruits factors that establish gene silencing or heterochromatin.^2,3^ This distinction highlights that the functional output of histone methylation critically depends on how chromatin-binding proteins recognize and bind their cognate histone substrates. Understanding the dynamics and mechanisms that regulate how histone-modifying enzymes localize in living cells is critical to defining how epigenetic states are established, maintained, and inherited.

In the fission yeast *Schizosaccharomyces pombe*, Clr4 is the only methyltransferase that catalyzes H3K9me, which subsequently leads to silencing at pericentromeric repeats, telomeres, and the mating-type locus.^4^ Conceptually, this process involves two phases: nucleation and spreading. During nucleation, sequence-specific recruitment factors (such as RNAi pathway proteins and DNA-binding proteins) recruit Clr4 to a defined genomic locus.^5^ Once established, H3K9me spreads to encompass large portions of the genome spanning dozens of kilobases of DNA. The coordination between two key structural domains within Clr4 is primarily responsible for nucleation and spreading: (1) the C-terminal SET domain catalyzes H3K9me (“writing”), and (2) the N-terminal chromodomain (CD) recognizes and binds H3K9me (“reading”).^6^ Beyond these structural features, Clr4 activity is additionally regulated through its assembly along with four other proteins (Cul4, Rik1, Raf1, and Raf2) into a multisubunit Cryptic Loci Regulator Complex (CLRC) (**Figure 1A**).^7–9^ The Cul4 E3 ubiquitin ligase has two key monoubiquitination targets: histone H3 (H3K14) and Clr4 itself.^10–12^ Together, these modifications fine-tune Clr4 enzymatic activity and promote its subsequent turnover from chromatin.^13^

**Figure 1.**
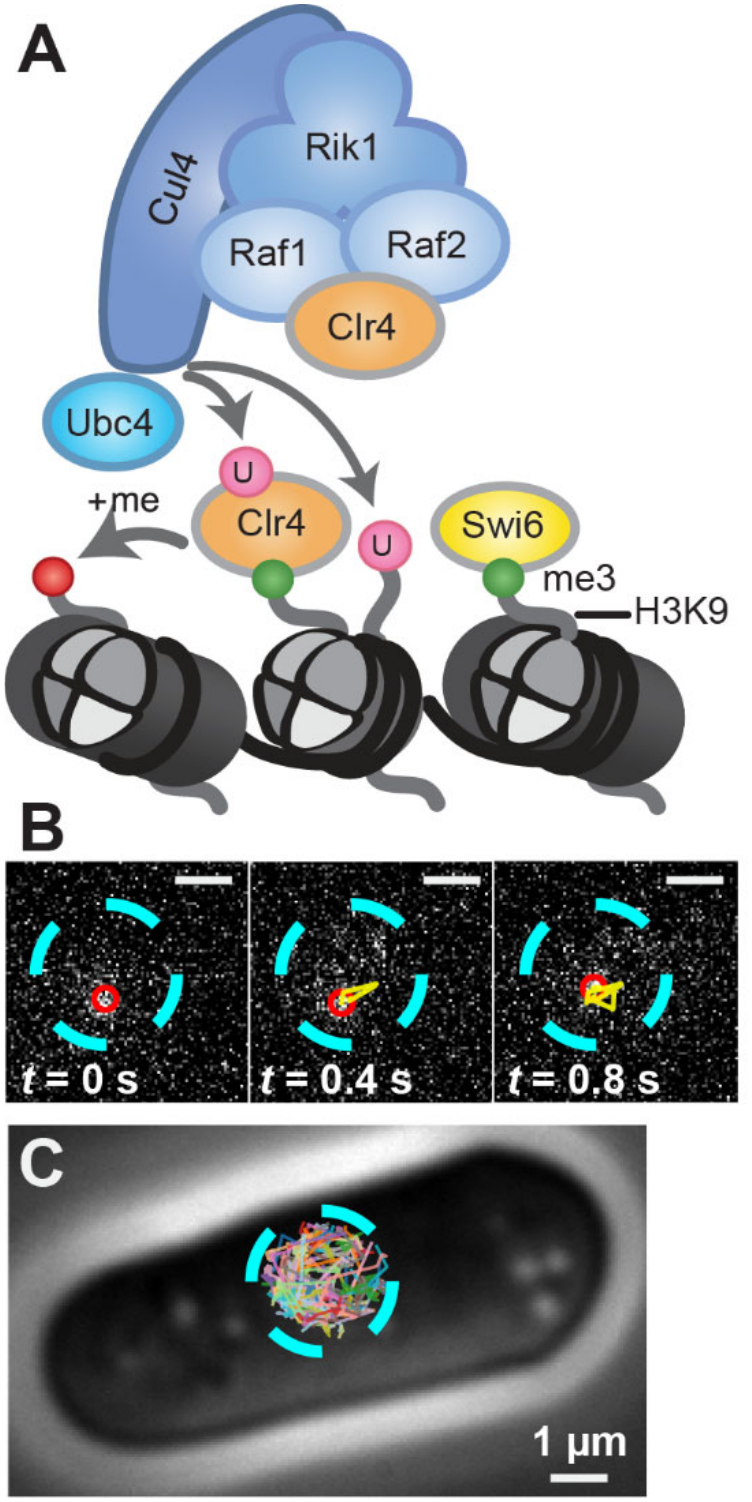
Single-molecule tracking of PAmCherry-Clr4 in living *S. pombe* cells. **(A)** Schematic of the multisubunit Clr4 methyltransferase complex (CLRC). Clr4-mediated H3K9 methylation depends on its enzymatic SET domain and its incorporation into a multisubunit complex consisting of Raf1, Raf2, Rik1, and Cul4 (an E3 ubiquitin ligase). Its enzymatic activity is additionally influenced by the ubiquitination of specific lysine residues within Clr4 itself and of its histone substrate (H3K14). Ubc4 is an E2 enzyme that promotes Clr4 ubiquitination. **(B)** Image of a single photoactivated PAmCherry-Clr4 molecule in the nucleus of an *S. pombe* cell at three time points. Red circle: molecule position; yellow line: Swi6-Clr4 trajectory from photoactivation until the current frame; cyan circle: approximate nucleus position. **_(C)_** Compilation of all single-molecule trajectories (false colors) acquired in the same *S. pombe* cell as in (B). Each trajectory is acquired after a new photoactivation cycle. Cyan circle: approximate nucleus position.

Despite our molecular understanding of the mechanisms that regulate Clr4 function, a fundamental problem remains unresolved. The *S. pombe* genome contains tens of thousands of nucleosomes, yet only a small fraction (∼2%) are within heterochromatic domains marked by H3K9me.^14^ After sequence-specific establishment, how does Clr4 efficiently discover and bind to these rare H3K9me sites? One possibility is that the Clr4 structural domains (SET and CD) are necessary and sufficient for chromatin localization. However, a second possibility has emerged, primarily based on *in vitro* studies: HP1 proteins, such as Swi6 in *S. pombe*, which bind H3K9me, may form phase-separated condensates that can trap Clr4, thus creating a microenvironment that sequesters the enzyme along with its substrate.^15^ In this second scenario, rather than searching the entire nuclear volume, Clr4 would partition into condensates with a dramatically elevated local concentration of H3K9me-marked nucleosomes. However, whether Swi6 condensates can in fact sequester Clr4 in cells remains untested. More broadly, how the native chromatin context, which involves chromatin being folded into higher-order structures with diverse post-translational histone modifications, influences Clr4 dynamics and affects its target search kinetics remains an open question.

Addressing these questions requires measurements of protein dynamics and intermolecular interactions between Clr4 and its binding partners in living cells in real time. Conventional approaches, such as co-immunoprecipitation to detect protein-protein interactions and chromatin immunoprecipitation to detect protein localization, capture interactions *in vitro* or after fixation and may therefore not reflect the true behavior of proteins in their native cellular environment. Single-molecule tracking in living cells overcomes these limitations by providing a direct readout of the dynamics of Clr4 and its interactions with chromatin.^15,16^ Live-cell microscopy is a non-invasive, real-time technology, and in vivo single-molecule fluorescence imaging attains nanometer-scale information about subcellular positions and millisecond-scale motions. Moreover, because binding events and biochemical interactions such as electrostatic interactions manifest as changes in the apparent diffusion rate, single-molecule tracking provides a sensitive readout of these interactions in their native cellular context.^17–21^

Our previous studies identified the biochemical properties of proteins based on the detection of phenotypes in single-molecule tracking: since intermolecular interactions decrease molecular mobility and increase binding times, mutations that affect biochemical interactions give rise to altered single-molecule dynamics. We measured the *in vivo* dynamics of the *S. pombe* HP1 proteins, Swi6 and Chp2, as well as the demethylase Epe1, the chromatin remodeler Mit1, and the deacetylase Clr3 in fission yeast.^16,22^ Here, we extend this approach to understand how the histone methyltransferase Clr4 detects and interacts with its heterochromatin substrate based on measuring the single-molecule dynamics of PAmCherry-Clr4 (**Figure 1B,C**). By tracking different Clr4 mutants within the *S. pombe* nucleus, we determined that the PAmCherry-Clr4 mobility is robust to changes in H3K9 methylation levels, chromatin structure, and Clr4 ubiquitination activity. Instead, Clr4 dynamics are driven by its primary structural domains, independent of protein sequestration mechanisms such as HP1-dependent condensates. Our results elucidate how Clr4 searches for and interacts with its heterochromatin substrate, providing insight into the dynamics that are critical for the establishment and maintenance of epigenetic states.

## Materials and Methods

### Plasmids, strains, and primers

*S. pombe* strains were constructed using a PCR-based gene targeting approach.^23^ The strains with PAmCherry fluorescent tags were created by constructing pDual vectors with the specified *nmt* promoter and the protein of interest (**Table S1**).^24^

### Sample preparation and single-molecule live-cell imaging in *S. pombe*

All yeast strains were grown on yeast extract with supplements and agar (YEA) plates for about 2 days at 32 °C. Afterward, individual colonies were inoculated in a standard YES medium (US Biological, cat. Y2060) containing the full set of yeast amino acids and incubated overnight at 32 °C with shaking. For strains expressing CLRC complex proteins from an *nmt1*, *nmt41*, or *nmt81* promoter, the fluorescence background was reduced by diluting the seed culture into Edinburgh Minimal Medium with Supplements and without Casamino acids (EMMC) medium (Formedium, cat. PMD0402) containing the full set of yeast amino acids, followed by incubation at 30 °C with shaking to reach an OD600 ∼0.5. To maintain cells in the log phase and prevent nuclear vacuole or spore formation, the culture was maintained at OD600 ∼0.5 for 2 days, with dilutions performed at ∼12-hour time intervals for YES medium cultures or ∼24-hour time intervals for EMMC medium cultures.

Samples for imaging were prepared by pipetting 1.5 – 2.5 µL of concentrated cells onto a pad of 1 – 2% agarose prepared in EMMC medium (for fluorescence background reduction) and then covered with an argon plasma-etched glass coverslip. Each agarose pad sample was imaged for less than an hour at room temperature to prevent the agarose from drying out. Samples were imaged in an Olympus IX71 epifluorescence microscope with a 100× 1.40 NA oil-immersion objective. The fluorescent background was decreased by bleaching preactivated PAmCherry molecules with exposure to 488-nm light (Coherent Sapphire 488, at a power density of 320 W/cm^2^) for 40 – 60 s and then exposure to 561-nm light (Coherent Sapphire 561-50, at a power density of 315 W/cm^2^) for 40 – 60 s. 50 – 100-ms activation pulses (2 – 5 W/cm^2^) of a 406-nm laser (Coherent Cube 405-100) were used for photoactivation, and a 561-nm laser (Coherent-Sapphire 561-50) was used for fluorescence excitation (520 W/cm^2^). The fluorescence emission was filtered to eliminate the 561-nm excitation source with a dual-band dichroic mirror and filter (CHROMA ZT488/561rpc and ZET488/561m-TRF, respectively) and imaged with a 40-ms exposure time per frame on a 512 × 512-pixel Photometrics Evolve electron-multiplying charge-coupled device (EMCCD) camera.

### Continuous imaging single-molecule tracking analysis

Single PAmCherry-Clr4 molecule positions were localized and tracked using the SMALL-LABS software (**Figure 1B,C**; **Supplementary Methods**, **Movie S1**).^25^ The nucleus of each cell was identified based on the autofluorescence outside the nucleus during the 488-nm bleaching step, and only signals within the nuclear region were included in the analysis. Only single-molecule trajectories with a minimum of 4 steps were used for trajectory analysis.

To determine the apparent diffusion coefficient, *D_app_*, of each trajectory, a modified diffusion model was applied to the mean square displacement, *MSD*, as a function of time lag, *τ*, over the range 40 ≤ *τ* ≤ 80 ms.^26^ This modified model accounts for the localization precision, *σ*:

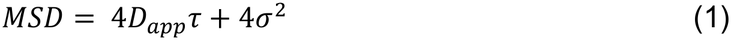

Fitting to an equation that explicitly accounts for motion blur based on the localization uncertainty and the finite camera exposure time did not change the results (**Eq. S1; Figure S4**).^27^ Fits were retained only if *R*^2^ ≥ 0.75. The logarithmic distribution histograms of *D_app_*were subsequently fit using a two-state Gaussian mixture model (GMM) to determine *D_app_* and the associated weight fraction for the slow and fast diffusive populations, respectively. We chose the two-component GMM because it allowed a reasonable physical interpretation in the cells. To calculate the 95% confidence intervals for the weight fractions and the mean *D_app_*, bootstrapping was performed by resampling 100% of the data across 10000 iterations. This method does not account for our hierarchical data collection methods and underestimates the true 95% confidence intervals. Numerical values are shown in **Figure S6** and **Tables S3** and **S4**.

Using tracks simulated with the built-in function s*imulate_tracks.sim_FOV* in exTrack,^28^ we validated that our methodology accurately measures the diffusion coefficients and population fractions of diffusing molecules. Two populations of molecules were simulated with *D* values and fractions taken from the GMM fit to our wild-type Clr4 data. To accurately simulate photobleaching, we fit the measured photobleaching rate for wild-type PAmCherry-Clr4 to a single-exponential decay function (**Figure S7**). The fit is good up to 20 imaging frames (0.8 s). Accordingly, we simulated tracks ranging from 1 to 20 frames long.

Overall, simulations largely reproduced the experimentally observed diffusion-coefficient peaks, with only a minor shift in the slow population, but underestimated the fraction of fast-moving molecules in the shortest track lengths, likely because fast-moving molecules are more likely to move out of focus and be lost from tracking. This detection bias is inherent to the single-molecule method but should not affect comparisons between strains because the same tracking limitations apply to all samples. Varying the minimum trajectory length had only a small effect on the calculated *D* (**Figure S5**).

### Anisotropy analysis

Datasets were pre-filtered to remove slow-moving (“fully bound”) trajectories (*D_app_* ≤ 0.1 μm^2^/s), as their inclusion could bias the angular distribution. The angle between each pair of consecutive displacements was calculated, and the anisotropy was assessed by comparing the fraction of angles within the range 180° ± 30° against those within 0° ± 30° across multiple length scales, calculated as:

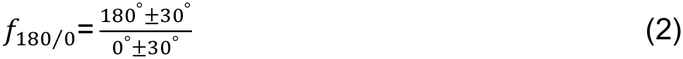

The anisotropy was plotted as a function of the mean of the two displacements forming the angle.^29^ To calculate 95% confidence intervals for the fold anisotropy, bootstrapping was performed by resampling 100% of the data across 10000 iterations. We confirmed that anisotropy values are not an artifact of imaging molecules at displacements near the localization precision by comparing anisotropy between observed and simulated tracks. In contrast to our experimental data, the simulated data has an almost perfectly isotropic distribution of steps (**Figure S8**).

### Slow imaging single-molecule tracking

Single-molecule trajectories were acquired using a 150-ms integration time with a 200-ms delay between consecutive frames. This imaging strategy blurs out PAmCherry-Clr4 molecules that remain mobile for a substantial portion of the frame while also enabling the use of reduced laser power (212 W/cm^2^) to limit photobleaching, resulting in longer trajectories.

Molecule positions were localized and tracked using the Trackmate software.^30^ The weight fractions of molecules in the slow- and fast-diffusing populations (*f*_sb_ and *f*_tb_, respectively) were determined by fitting a two-population diffusion model to the cumulative distribution function (CDF) of the squared displacements as previously described:^31^

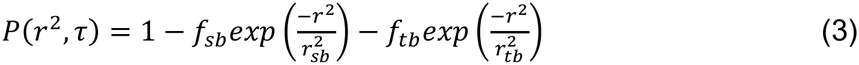

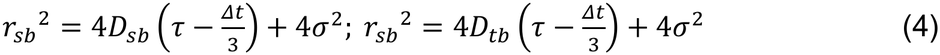

In these equations, *r* ^2^ are the experimentally observed squared displacements, *D*_sb_ and *D*_tb_ are the diffusion coefficients of the slow- and fast-diffusing populations, respectively, *τ* = 350 ms is the total time per frame, and Δ*t* = 150 ms is the integration time. The localization precision, *σ* = 30 nm, was estimated based on previous experiments at similar imaging conditions.^16^ Equations (3) and (4) were fit to experimental data by least-squares curve fitting using the Python function *minimize* from the SciPy library.^32^

The stable binding residence time (*τ_sb_*) for each strain was extracted from the survival probability curves, which were compiled from single-molecule trajectories for each strain. These curves were fit to a double exponential decay function:

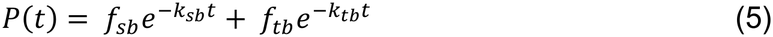

In this model, the off-rates for stable and transient binding events are the slower *k_sb_* and the faster *k_tb_*, respectively, while *f_sb_* and *f_tb_* represent their relative fractions (*f_sb_* + *f_tb_* = 1). The weights *f_sb_* and *f_tb_*were set to the weight fractions of molecules in the slow- and fast-diffusing populations previously calculated from the CDF. Since the observed *k_sb_* is influenced by photobleaching and chromatin dynamics, bias correction was performed by subtracting the *k_sb_* obtained for the nominally immobile H2B-PAmCherry from that of PAmCherry-Clr4 across different strains.^33,34^ The corrected *τ_sb_* was calculated as the reciprocal of the adjusted *k_sb_* (**Table S5**). 95% confidence intervals were calculated by fitting a double-exponential decay function to 100% of the data bootstrapped 10000 times.

Fractions were fixed to values from a CDF because the double-exponential model showed extremely high variability otherwise. Residence times were calculated with CDF-fixed amplitudes (**Figure S9A**) and amplitudes that were allowed to vary (**Figure S9B**). We cannot definitively state that residence times remain constant between methods because the confidence intervals become extremely large when the amplitudes can vary.

### Single-molecule RNA fluorescence in situ hybridization (smFISH)

*S. pombe* cells were grown to mid-log phase in Edinburgh minimal medium with supplements (EMMC), and a total of 12 OD of cells were harvested. Based on earlier reports,^35–37^ the cells were fixed using 3% formaldehyde at room temperature on a rotator for 20 min and washed 3× in 1.2 M Sorbitol. Cells were then spheroplasted in 1 M sorbitol, 50 mM Tris pH 7.5, 5 mM *β*-mercaptoethanol, and 2 mg/mL Zymolyase at 37 °C for about 45 min. The reaction was considered complete when all the cells appeared translucent upon addition of 1% SDS. Cells were then resuspended in 70% ethanol and incubated at room temperature for 4 hr, followed by resuspension in a formamide wash buffer (FWB; 10% Formamide, 2× SSC) for 20 min at room temperature.

For RNA FISH probe hybridization, cells were resuspended in 100 µL hybridization buffer (10% dextran sulfate, 1 mg/mL *E. coli* tRNA, 2 mM vanadyl ribonucleoside complex (VRC), 0.2 mg/mL BSA, 2× SSC, 10% formamide). 10 µL of 200 µM VRC and 2 µL of 2.5 µM probe were added to the sample, and the mixture was incubated for 16 hr at 30 °C in the dark with shaking at 75 rpm. Following hybridization, cells were washed 2× in FWB for 30 min each at 30 °C, resuspended in FWB containing DAPI (1 μg/mL final concentration), and incubated at 30 °C for 30 min. Finally, the cells were resuspended in imaging buffer (2× SSC, 50 mM Tris pH 8.0) and stored at 4 °C until imaging. The RNA FISH probes used in this study were designed with the Stellaris Probe Designer and purchased from LGC Biosearch Technologies. The probe set comprises 48 20-nt oligonucleotides covering the full length of the *clr4* transcript, each conjugated to Quasar 670.

Fluorescence images were acquired on a widefield epifluorescence microscope (Nikon Ti2) equipped with a 100×/1.49 NA oil-immersion objective and imaged with a DAPI and Cy5 filter set. *Z*-stacks of 19 optical sections were recorded at 0.2-µm spacing. Exposure times and acquisition settings were kept constant for all samples. Cell segmentation was performed in Python using Cellpose^38,39^ after training a model with a manually labelled dataset of cells. Nuclear segmentation and spot analysis were performed using customized scripts built on Big-FISH^40^. smFISH images were analyzed on 2D maximum-intensity projections of the *z*-stacks. For each dataset, the intensity threshold and spot-size parameters for spot detection were chosen by visual inspection in representative images and kept constant across all images in that dataset.

## Results

### PAmCherry-Clr4 is functional to establish silencing and can be expressed at optimal levels for single-molecule imaging

To track individual Clr4 molecules in *S. pombe* cells, we fused Clr4 to a photoactivatable PAmCherry protein.^41^ We expressed *pamCherry-clr4* under the control of three thiamine-repressible promoters: *nmt1*, *nmt41*, or *nmt81* in *clr4Δ* cells (**Figure 2A**); these promoters are actively transcribed when thiamine is absent.^42^ The promoters exhibit graded expression, and we classify them as high (*nmt1*), medium (*nmt41*), and low (*nmt81*) expression promoters. Through western blots, we confirmed that the *nmt1* promoter exhibits the highest level of PAmCherry-Clr4 expression, the *nmt41* promoter produces moderate levels of expression, and the *nmt81* promoter yields the lowest expression levels (**Figure S10**). We compared *clr4* transcript levels expressed from different promoters using single-molecule FISH, and we determined that the *nmt81*, *nmt41*, and *nmt1* promoters increase transcript levels 2-, 3-, and 19-fold relative to the native *clr4* promoter, respectively (**Figure 2B,C,D**). To evaluate how Clr4 expression levels and the PAmCherry fusion affect heterochromatin function, we tested the silencing of a *ura4+* reporter gene inserted at the mating type locus, which is an endogenous site of heterochromatin formation in *S. pombe* (*Kint2::ura4+*).^43^ PAmCherry-Clr4 expression using the *nmt1* promoter disrupted silencing, whereas the *nmt41* and *nmt81* promoters preserved *ura4+* reporter silencing, as indicated by growth of cells on FOA-containing medium. However, only expression from the *nmt41* promoter additionally led to reduced viability of cells on −URA medium, resembling what we typically observe in wild-type control cells (**Figure 2E**). These results suggest that a high-strength promoter (*nmt1*) disrupts silencing and a low-strength promoter (*nmt81*) produces an unstable silencing phenotype. The *nmt41* promoter therefore provides an optimal expression level that preserves wild-type heterochromatin function, and we used this strain for all subsequent experiments.

**Figure 2.**
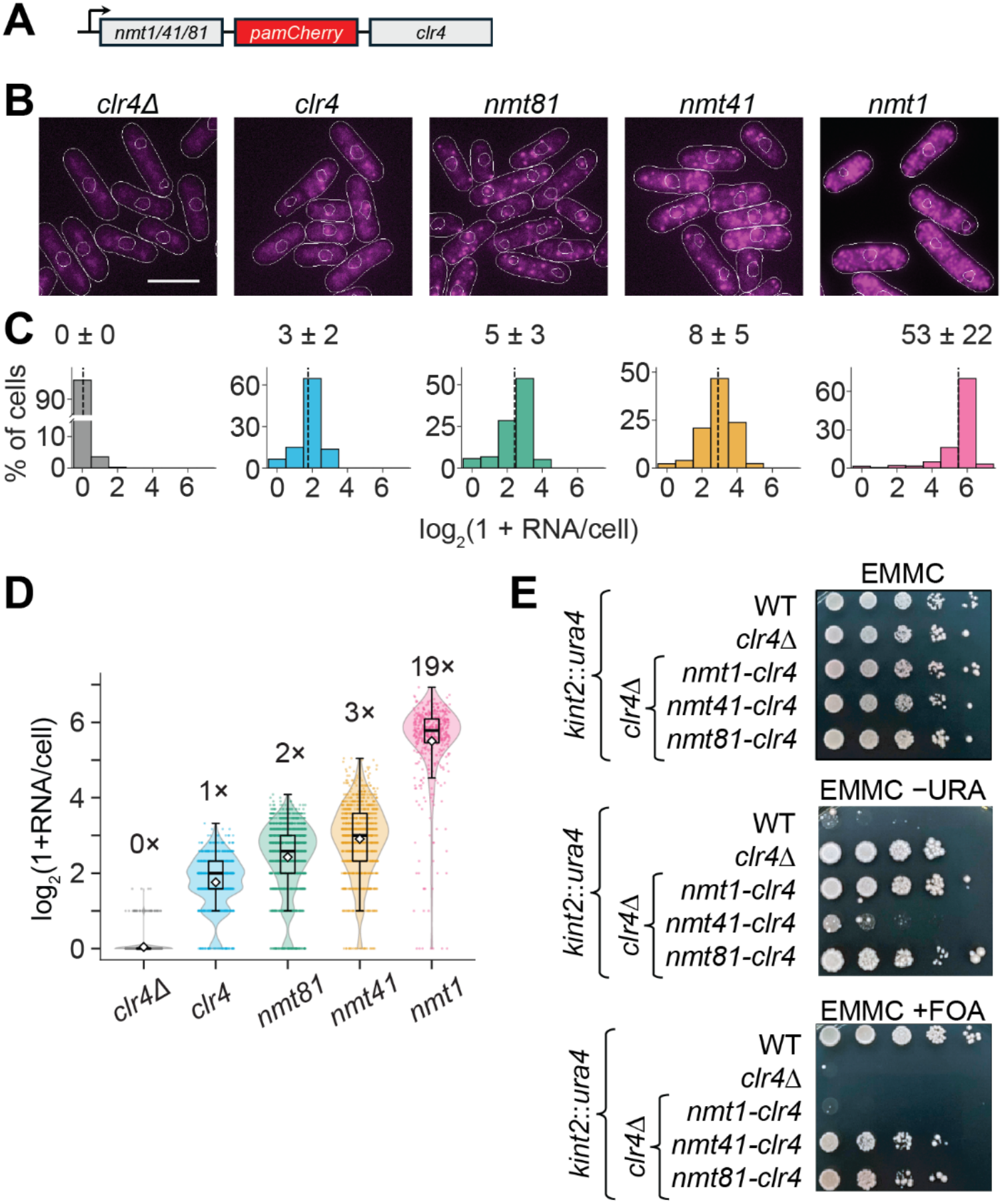
Validation of PAmCherry-Clr4 transcription and its effect on heterochromatin establishment at the mating type locus. **(A)** PAmCherry is fused to the N-terminus of Clr4 and expressed ectopically using an inducible promoter: *nmt1, nmt41,* or *nmt81* (high, medium, or low expression levels, respectively). **(B)** Single-molecule FISH images of *clr4* transcripts expressed from different promoters, including the native *clr4* promoter and the inducible promoters in (A). Each bright pink spot represents a single transcript. White outlines: cell and nucleus boundaries. Scale bar: 10 µm. **(C)** Distribution of RNA spots per cell. A pseudo-count of 1 was added so that cells with zero transcripts appear at 0. Dashed lines mark the mean of log₂(1 + RNA/cell), and the mean ± SD of the actual RNA counts is given above each histogram (*clr4Δ*: 0.04 ± 0.22 RNA/cell, *n* = 1165 cells; *clr4* promoter: 2.7 ± 1.6, *n* = 765; *nmt81*: 5.3 ± 3.0, *n* = 1550; *nmt41*: 8.0 ± 5.3, *n* = 1006; *nmt1*: 52.7 ± 21.5, *n* = 503). **(D)** Distribution of RNA spots per cell for the same samples as in (C). Each dot is one cell. Violins: density, boxes: medians and quartiles; whiskers extend to the lowest and highest cells within 1.5× of the interquartile range. White diamonds: mean of log₂(1 + RNA/cell). Fold change of each sample’s mean RNA count per cell relative to *clr4* promoter is indicated above each dataset. **(E)** Silencing assay using a *ura4+* reporter inserted at the mat locus (*Kint2::ura4*). 10-fold serial dilutions of cells expressing PAmCherry-Clr4 from different *nmt* promoters were plated on EMMC, EMMC +FOA, and EMM −URA plates.

### Measurements of Clr4 dynamics identify slow-moving chromatin-bound molecules and fast-moving unbound molecules

We tracked single PAmCherry-Clr4 molecules in real time with continuous imaging and 40-ms/frame integration times in otherwise wild-type (WT) cells (*clr4+*) (**Figure 1B,C**; **Movie S1**). The distribution of the single-trajectory diffusion coefficients indicates two populations: we found that 46% of Clr4 molecules diffuse slowly (average *D*_slow_ = 0.030 μm²/s) and the remaining 54% of Clr4 molecules diffuse quickly (average *D*_fast_ = 0.45 μm²/s) (**Table S2, Figure 3A**). Based on our previous measurements of HP1 proteins, we interpret the slow mobility state as chromatin-bound Clr4 and the fast mobility state as corresponding to mostly diffusing, unbound Clr4 molecules.^16,22^

**Figure 3.**
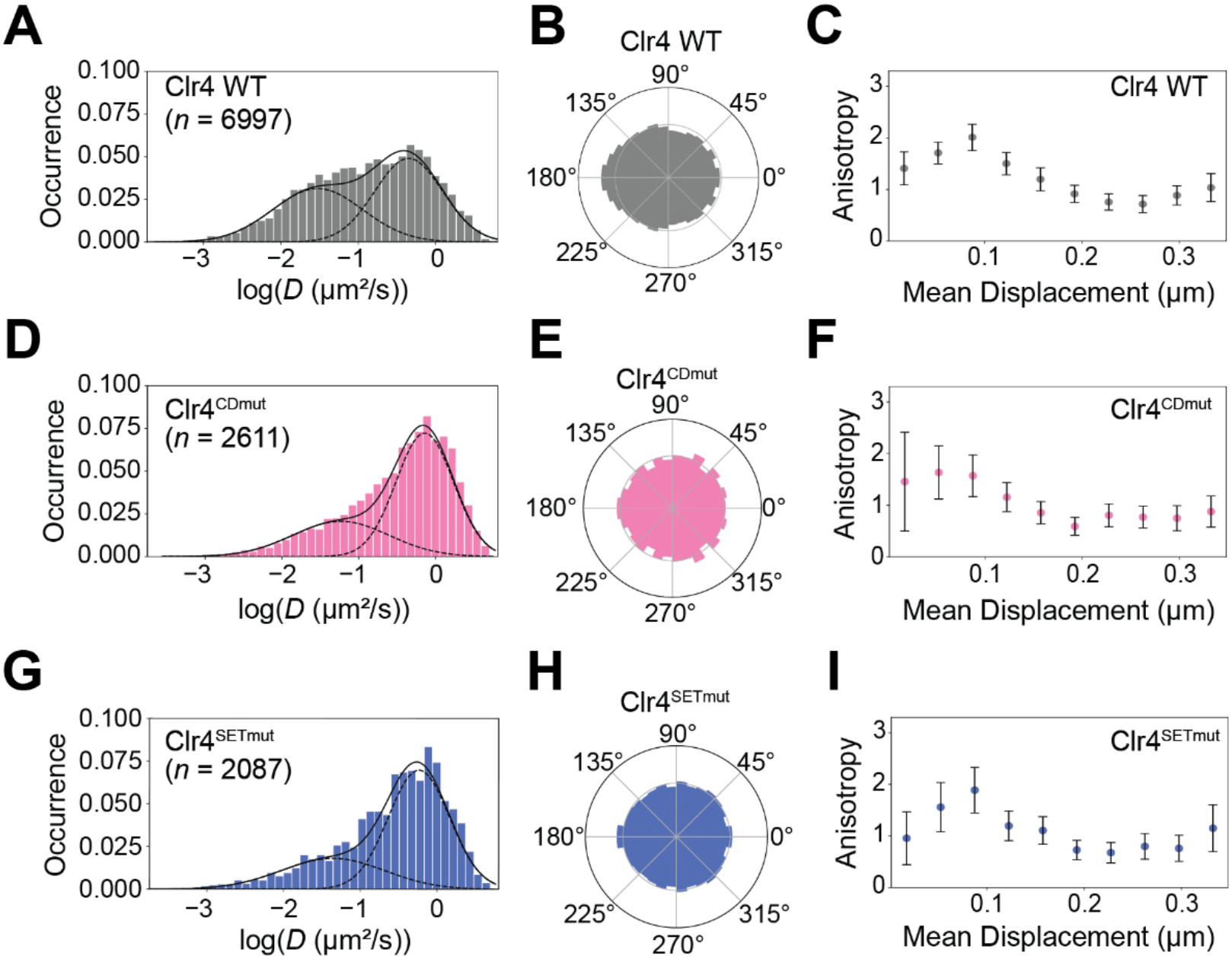
PAmCherry-Clr4 single-molecule dynamics for wild-type Clr4, Clr4 with a chromodomain mutation (Clr4^CDmut^), and Clr4 with a SET domain mutation (Clr4^SETmut^). (A,D,G) Diffusion coefficient distribution of PAmCherry-Clr4 trajectories in each indicated *S. pombe* strain. Distributions are fit to a two-term Gaussian mixture model (dashed lines) to estimate the fast and slow diffusion populations (Table S2). *n* denotes the number of trajectories measured in each condition. **(B,E,H)** Polar histograms of angles within the mobile single-molecule trajectories with *D_app_* ≥ 0.1 μm^2^/s in panels A,D,G. **(C,F,I)** Anisotropic diffusion measurement (*f*_180/0_) of the faster population of diffusing PAmCherry-Clr4 molecules in each strain (*D*_app_ ≥ 0.1 μm^2^/s). Error bars indicate the 95% confidence intervals from bootstrapping 100% of the data across 10000 iterations.

We also calculated the angles between consecutive displacements within the selected mobile single-molecule trajectories (*D*_app_ ≥ 0.1 μm^2^/s) to determine the angular distribution for the motion of fast-moving (biologically interpreted as unbound) PAmCherry-Clr4 molecules during target search. This distribution would be uniform for random (Brownian) diffusion (fold anisotropy, *f*_180/0_ = 1.0). Instead, we measured mildly anisotropic Clr4 motion (*f*_180/0_ = 1.40) at very small displacements (∼0.018 μm), indicating a bias toward reversing direction (180° turns) (**Figure 3B**). This anisotropy rises to a peak of ∼2 at intermediate displacements (0.07 – 0.1 μm), and it becomes ∼1.0 (no anisotropy) for larger displacements (> 0.175 μm). This analysis implies that the motion of Clr4 molecules during the search for their cognate binding site is spatially constrained and distance-dependent relative to sites of heterochromatin formation (**Figure 3C**, **Figure S1**).

### The Clr4 chromodomain and SET domain determine the fraction of slow-moving chromatin-bound Clr4 molecules

We examined how the chromodomain (CD) affects the binding of Clr4 to its cognate histone substrate. A single amino acid change (W31G) within the Clr4 chromodomain disrupts H3K9me recognition, binding, and spreading.^6^ We measured the dynamics of this PAmCherry-Clr4^CDmut^ expressed from the *nmt41* promoter in cells where the endogenous Clr4 copy was deleted (*clr4Δ*). Hence, the only copy of Clr4 that these cells express is a CD-defective version. We expected to observe less H3K9me binding by PAmCherry-Clr4^CDmut^ relative to PAmCherry-Clr4, and indeed, the weight fraction of the slow population decreased from 46% to 33% (**Figure 3D, Table S2**). We also measured a greater diffusion coefficient for the fast mobility state, consistent with the fast PAmCherry-Clr4^CDmut^ molecules being predominantly unbound (*D*_fast_ = 0.72 μm²/s for Clr4^CDmut^ vs. *D*_fast_ = 0.45 μm²/s for wild-type Clr4) (**Figure 3D, Table S2**).

Notably, the slow diffusion coefficient also increased (*D*_slow_ = 0.06 μm²/s for Clr4^CDmut^ vs 0.030 μm²/s for wild-type Clr4). Since Clr4^CDmut^ cannot specifically bind H3K9me, the residual slow population likely represents transient, nonspecific interactions rather than binding to its cognate histone substrate, and indeed, the Clr4^CDmut^ dynamics are more isotropic (**Figure 3E**), including at intermediate displacements (**Figure 3F, Figure S1**). The anisotropy profile of Clr4^CDmut^ is consistent with transient interactions, which we attribute to a chromatin-sampling state, in which Clr4 binds to and searches for its cognate ligand.

To determine how catalytic activity affects Clr4 dynamics, we introduced two amino acid substitutions (H410L C412A) that attenuate H3K9 methyltransferase activity (Clr4^SETmut^).^44^ We measured PAmCherry-Clr4^SETmut^ dynamics when expressed from the *nmt41* promoter in cells lacking endogenous Clr4. Hence, all Clr4 binding events in this strain background are interactions between Clr4 and unmethylated histone substrates. Given the complete absence of H3K9me in this background, we expected to measure a loss of chromatin binding upon tracking PAmCherry-Clr4^SETmut^, and indeed, the weight fraction of the slow population decreased from 46% to 31% (**Figure 3G, Table S2**). As with the Clr4^CDmut^, we interpret the residual slow population as transient, chromatin-sampling interactions rather than specific binding to H3K9me chromatin. We noted that the fast and slow Clr4^SETmut^ diffusion coefficients are both intermediate in value between the wild-type Clr4 diffusion and the Clr4^CDmut^ dynamics (**Figure 3G, Table S2**). These results suggest that, although the SET domain mutation disrupts H3K9me, the Clr4^SETmut^ protein retains its ability to sample histones in search of its cognate histone substrate. The anisotropy profile for PAmCherry-Clr4^SETmut^ is similar to that of PAmCherry-Clr4^CDmut^, including more isotropic behavior (more freely diffusive motion) (**Figure 3H,I, Figure S1**). This isotropy supports the interpretation that Clr4^SETmut^ molecules fail to recognize their targets and only transiently bind to nucleosome substrates.

Besides its structural domains. Clr4 activity is additionally regulated through its incorporation into the multisubunit Cryptic Loci Regulator Complex (CLRC) (**Figure 1A**). We expressed PAmCherry-Clr4 in cells in which the CLRC scaffold protein Rik1 was deleted, and found no change in the single-molecule diffusion coefficients or single-step anisotropies for Clr4 in this *rik1Δ* background relative to wild-type cells (**Figures S1 and S2G-I**), These results indicate the Clr4 dynamics we measured are independent of the incorporation of the protein into the multisubunit CLRC complex.

### Clr4 is recruited to heterochromatin independent of Swi6 condensates

HP1 family proteins bind to H3K9me and play conserved roles in heterochromatin establishment and maintenance. In *S. pombe,* the primary HP1 protein is Swi6, which has multiple functions during heterochromatin assembly. Based on *in vitro* data, one possibility is that HP1 proteins can sequester or trap enzymes such as Clr4 at sites of H3K9me (**Figure 4A**). To determine whether Swi6 condensates regulate Clr4 binding, we expressed PAmCherry-Clr4 from the *nmt41* promoter in cells in which Swi6 is deleted (*swi6Δ*). Comparing wild-type and *swi6Δ* strains, we observed no significant change in either the diffusion coefficients or the weight fractions of the slow and fast Clr4 populations, indicating that Clr4 must be recruited to heterochromatin through its structural domains (the CD and SET domains) but not through any interactions involving Swi6 (**Figure 4B,E, Table S2**). The single-molecule anisotropy within the PAmCherry-Clr4 tracks in *swi6Δ* cells is also similar to the wild-type background, which indicates that, even in the absence of Swi6, Clr4 motion continues to be constrained at intermediate distances from heterochromatin (**Figure 4C-D, F-G, Figure S1**).

**Figure 4.**
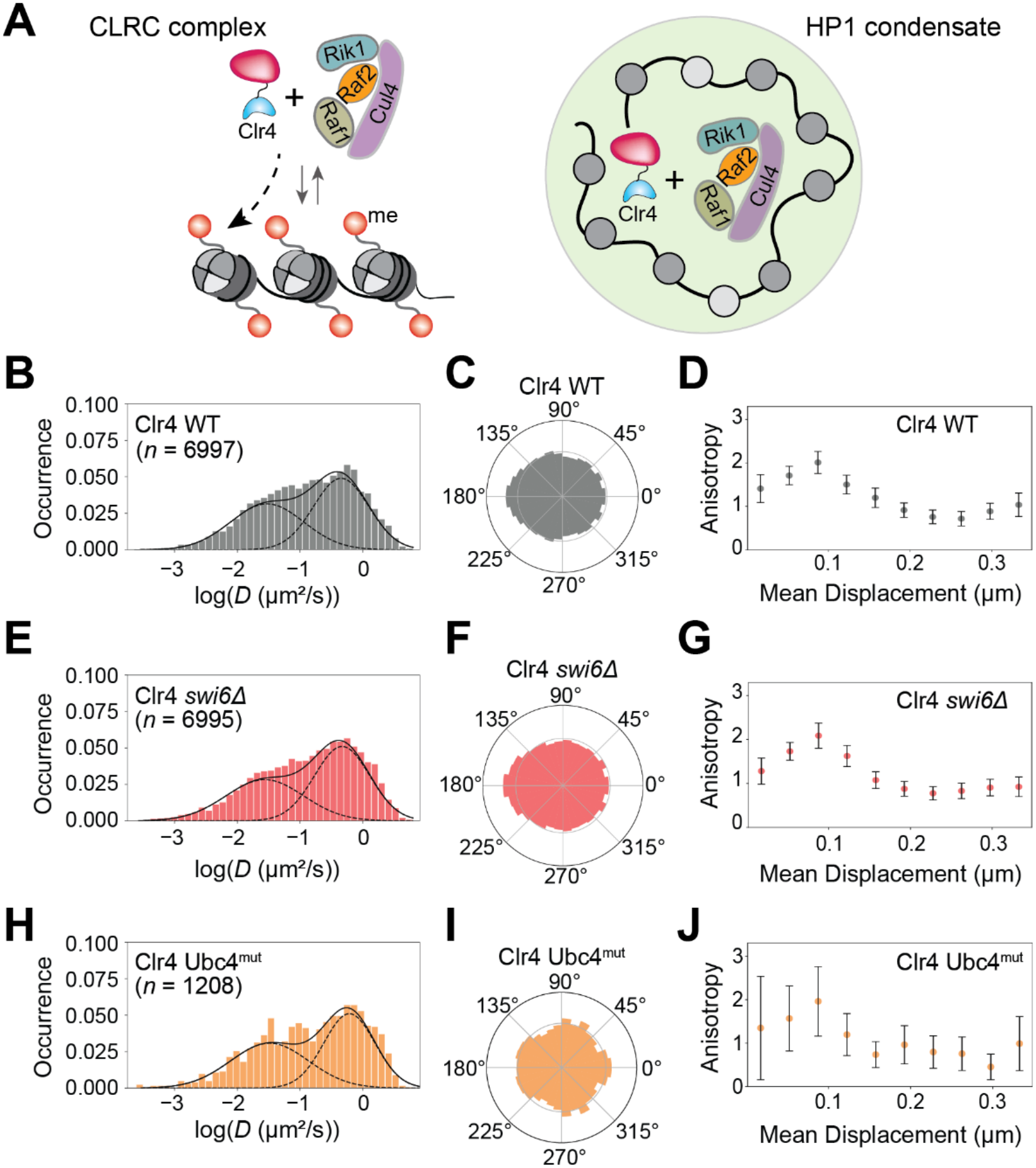
PAmCherry-Clr4 single-molecule dynamics in (B – D) WT *S. pombe* and in *S. pombe* mutants: (E – G) Swi6 deletion and (H – J) Ubc4 mutant. **(A)** Schematic of the intrinsic binding model (Clr4 dynamics in cells are driven by direct chromatin binding and unbinding) and the condensate-based trapping model (HP1 proteins sequester enzymes such as Clr4 at sites of H3K9me). **(B,E,H)** Diffusion coefficient distribution of *n* PAmCherry-Clr4 trajectories in each indicated *S. pombe* strain. Distributions are fit to a two-term Gaussian mixture model (dashed lines) to estimate the fast and slow diffusion populations (Table S1). *n* denotes the number of trajectories measured in each condition. **(C,F,I)** Polar histograms of angles within the mobile single-molecule trajectories with *D_app_* ≥ 0.1 μm^2^/s in panels B,E,H. **(D,G,J)** Anisotropic diffusion measurement (*f*_180/0_) of the faster population of diffusing PAmCherry-Clr4 molecules in each strain (*D*_app_ ≥ 0.1 μm^2^/s). Error bars indicate the 95% confidence intervals from bootstrapping 100% of the data across 1000 iterations.

### The ubiquitination of Clr4 leads to increased chromatin turnover at fast time scales

A second mechanism that has been proposed to contribute to Clr4 dynamics involves the ubiquitination of Clr4 within its intrinsically disordered hinge region through the activity of an E2 ubiquitin-conjugating enzyme, Ubc4 (**Figure 1A**). A Ubc4 mutant, Ubc4 G48D, reduces Clr4 ubiquitination, and ChIP-seq data indicate that the absence of ubiquitination leads to increased Clr4 chromatin occupancy.^13^ To test how ubiquitination shapes Clr4 dynamics, we expressed PAmCherry-Clr4 in Ubc4 G48D cells. Compared to the wild-type Clr4 background, PAmCherry-Clr4 in this Ubc4^mut^ background has a similar slow state diffusion coefficient (*D*_slow_ = 0.033 μm²/s) and slow population fraction (48%) as in the WT background. In contrast to the current ChIP-seq data, our findings indicate that the interaction of Clr4 with its chromatin substrate does not depend on ubiquitination. However, the fast population of Clr4 diffuses more quickly in the Ubc4^mut^ background (*D*_fast_ = 0.6 μm²/s), suggesting that Clr4 is more completely unbound from chromatin (**Figure 4H, Table S2**). The anisotropy results also support this observation, indicating more isotropic motion for Clr4 in Ubc4^mut^ cells, which corresponds to its more complete unbinding from chromatin (**Figure 4I,J, Figure S1**). These results suggest that the lack of ubiquitination attenuates Clr4 interactions with chromatin as opposed to enhancing its binding affinity. This type of discrepancy between single-molecule binding data and genome analysis has also previously been observed in the case of the transcription factor Sp1,^45^ highlighting how measurements at faster timescales or crosslinking conditions produce deviations from chromatin binding measurements in living cells.

### Histone substrate mutations do not affect the Clr4 search dynamics

We measured the PAmCherry-Clr4 dynamics in a strain with the H3K9 lysine modified to arginine (H3K9R). This modification blocks H3K9me and prevents heterochromatin formation similar to *clr4Δ*, and the Clr4 CD loses its high-affinity docking in the H3K9R background.^46,47^ We therefore expected to measure a significant decrease in the population of slower Clr4 molecules in the H3K9R cells. However, the slow population is unchanged in the H3K9R background relative to wild-type cells (**Figure S2D**, **Table S2**). Still, Clr4 diffuses faster in the H3K9R background than in wild-type cells (*D*_slow_ = 0.040 μm²/s and *D*_fast_ = 0.64 μm²/s), indicating that Clr4 dynamics in wild-type cells reflect some specific H3K9 binding that is lost in the H3K9R mutant. Furthermore, the anisotropy profile of PAmCherry-Clr4 is generally unchanged in the H3K9R cells relative to the wild-type cells (**Figure S1**, **Figure S2E,F**). This result suggests that Clr4 can search and interact with its chromatin substrate, even when H3K9me catalysis is prevented.^48^ This finding indicates that the more significant changes in the Clr4^CDmut^ dynamics relative to those of wild-type Clr4 (**Figure 3D-F**, **Table S2**) are attributable to structural changes in how the CD domain interacts with the histone substrate rather than changes in substrate composition.

### The Clr4 residence times are independent of specific binding

Our 40-ms time-resolution single-molecule tracking captures millisecond-scale dynamics and transient interactions, while longer timescale measurements access slower binding and unbinding events that occur on the scale of several seconds. We therefore imaged with a 200-ms time lapse between each 150-ms imaging frame to reduce photobleaching and enable longer trajectory measurements that capture the stability of the chromatin-bound state of Clr4. With these imaging parameters, we measured the apparent PAmCherry-Clr4 residence times in each genetic background based on the distribution of track lengths (**Figure 5A**). Log-scale analysis of the survival curves indicated that some of the fast motion was still detected even with this slow imaging condition, so we fit each corrected survival curve to a biexponential decay function (**Figure 5A**) based on weights chosen from a fit to the cumulative distribution function (CDF) of the squared displacements (**Figure S3**). We corrected the rate, *k_sb_*, of the slower term for photobleaching by comparison to essentially immobile histones (H2B-PAmCherry; **Figure 5A**).^33^ The reciprocal of this corrected slow-binding rate is the residence time of Clr4 on chromatin in that strain (Methods, **Table S5**).^31^

**Figure 5.**
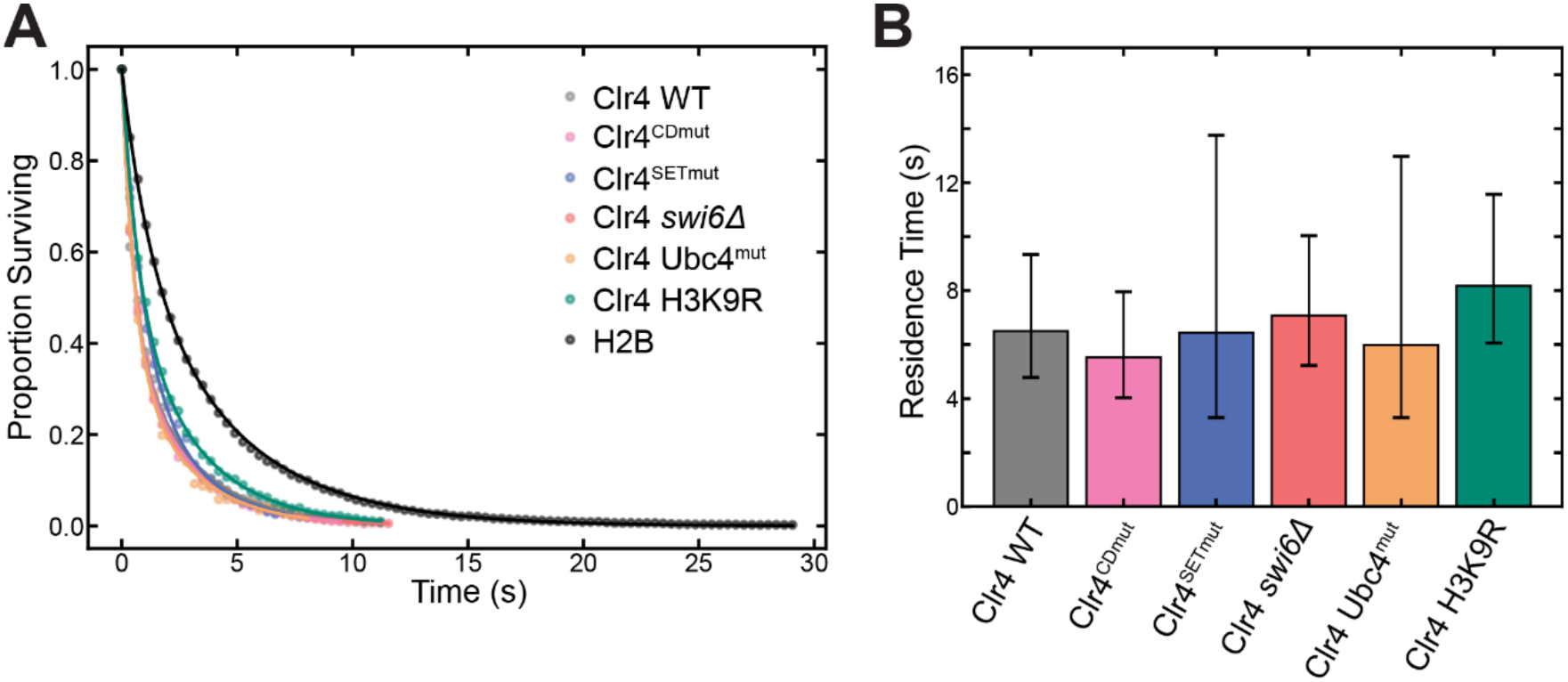
PAmCherry-Clr4 residence time calculation based on slow-frame time-lapse imaging. **(A)** The survival probability curve decay is fit to *P*(*t*) = *f_sb_e*^−k^*^sbt^* + *f_tb_e*^−k^*^tbt^*, where the weights, *f_sb_* and *f_tb_*, are determined from the CDF fits in Figure S3. **(B)** Each adjusted residence time is calculated from a fit to the corresponding decay curve in A and corrected based on the apparent decay of the nominally immobile H2B-PAmCherry. 95% confidence intervals were calculated by fitting a double-exponential decay function to 100% of the data bootstrapped 10000 times.

We found that PAmCherry-Clr4 has an average residence time of ∼6 s (**Figure 5B**). Surprisingly, this residence time was not significantly changed for any of the mutants measured (**Figure 5**). Overall, within the resolution of our assay, this measured residence time of 5 – 8 s is therefore independent of whether Clr4 can specifically bind to H3K9me (**Table S5**). Hence, the residence times primarily report on Clr4 binding to unmethylated histones, suggesting chromatin binding generally, rather than H3K9me binding specifically, drives our measurements of Clr4 dynamics.

## Discussion

In this study, we show that the Clr4 structural domains involved in H3K9me binding (chromodomain) and catalysis (SET domain) are the primary determinants of Clr4 dynamics in living *S. pombe*. None of the other perturbations that we investigated considerably affected the Clr4 dynamics: in all of these other experiments, we observed both slow, chromatin-associated PAmCherry-Clr4 and fast, freely diffusing PAmCherry-Clr4, and we measured only modest differences in the relative fractions and diffusion coefficients of these two populations (**Figures 3, 4, S2**). Previous studies that have measured CLRC complex dynamics by visualizing Raf2-Halo measured differences in how fast molecules move in wild-type cells versus a Raf1 overexpression background.^49^ Our measurements of Clr4 show that the enzyme samples the nucleus with different dynamics from proteins that are part of the CLRC scaffold. This robustness suggests that functional mutations that alter the *in vitro* Clr4 enzymatic activity and the H3K9me distribution in cells do not necessarily affect how Clr4 interacts with chromatin. Indeed, no genetic background fully eliminated the slow state, unlike what we previously measured in the case of HP1 proteins, for which the slow state is very sensitive to the presence of H3K9me.^16,22^ The persistence of a detectable slow population, even in the case of Clr4^CDmut^ and Clr4^SETmut^, suggests that Clr4 can still engage chromatin after structural disruptions via alternative binding modes that likely involve intrinsically disordered regions (IDRs).^48,50^ IDR sequences have several lysine and arginine residues, which have been implicated in binding non-specifically to nucleic acids and chromatin.^16,51^

Based on our data, we propose that Clr4 engages a di-nucleosome substrate in cells, with the CD binding one nucleosome and the SET domain binding a neighboring nucleosome in a configuration primed for read-write activity.^52^ Consistent with this model, we measured no change in Clr4 dynamics in H3K9R strains, in which the H3 histone is unmethylatable, which indicates that chromatin binding does not require a substrate that is amenable to methylation. Rather, the ability of both structural domains to bind the cognate histone substrates is the primary determinant of Clr4 chromatin binding kinetics *in vivo*. These observations also imply that, even in wild-type cells, our measurements primarily capture Clr4 interactions with unmethylated histone substrates and are not necessarily driven by heterochromatin-associated H3K9 methylation. This bias is likely because only ∼2% of all nucleosomes are methylated in *S. pombe* cells, and because Clr4 is promiscuous in its ability to bind to chromatin, unlike HP1 proteins, which localize at sites of H3K9 methylation with high specificity. This high level of chromatin sampling or non-specificity with which Clr4 interacts with chromatin may explain how Clr4 can sample H3K9me at novel genomic sites under stress conditions without an explicit requirement for recruitment by sequence-specific factors.^53,54^

In most genetic backgrounds, the mild anisotropy in the motion of Clr4 at short and intermediate displacements suggests that Clr4 does not sample chromatin just by free three-dimensional diffusion. Instead, the absence of anisotropy at longer displacements indicates a guided exploration mode in which Clr4 molecules repeatedly sample nearby chromatin before fully disengaging. This behavior supports a model where Clr4 periodically probes nucleosomes and other chromatin-associated complexes during its search, increasing the likelihood of finding proper H3K9 sites and promoting heterochromatin formation.

The primary role of the chromodomain and SET domain in governing Clr4 chromatin binding indicates that HP1 proteins do not drive Clr4 chromatin occupancy through sequestration.^13^ In support of a Clr4 intrinsic binding model that does not require other protein binding interactions, we find that deletion of the HP1 homolog Swi6 does not reduce the fraction of slow Clr4 molecules, nor does it alter the Clr4 diffusion coefficient or its residence time on chromatin under the conditions examined, although smaller or spatially restricted effects cannot be excluded.^22^

## Conclusion

Overall, our results demonstrate that Clr4 operates *in vivo* as a methyltransferase that robustly maintains a chromatin-bound population across a broad range of genetic perturbations while constantly exploring the nuclear environment. The Clr4 chromodomain and SET domain guide this exploration process, while incorporation of Clr4 into the CLRC complex or post-translational modifications that affect its enzymatic activity have no effect. Rather than sequestration-based mechanisms, we propose that Clr4 dynamics in cells are driven by direct chromatin binding and unbinding mediated by the CD and SET domains. Given their conservation across organisms, we expect that homologous enzymes such as the human Suv39h will share similar dynamic properties.^55^

## Study Limitations

In our single-molecule tracking measurements, trajectory lengths are constrained by photobleaching, blinking, and axial diffusion out of the focal plane. Because rapidly diffusing molecules can exit the focal plane faster than slow or bound molecules, fast-moving populations may be underrepresented. Together with localization uncertainty, motion blur, and finite frame rates, these factors limit the precision of absolute diffusion coefficients and state fractions, so our focus here is on a comparative analysis of how these factors change across different genetic backgrounds. Since heterochromatin comprises only ∼2% of the *S. pombe* genome, our nucleus-wide SMT measurements likely capture broad chromatin sampling alongside locus-specific silencing. Thus, observed mobility shifts may reflect either direct changes in Clr4 affinity or indirect alterations to the local chromatin microenvironment.

## Acknowledgments

We thank Danesh Moazed, Songtao Jia, and Rob Martiennsen for sharing *S. pombe* strains with us. Support for this work comes from National Science Foundation grant MCB-2602737 to KR and JSB, a National Science Foundation grant CHE-2403937 to JSB, an NIH award R35GM137832 to KR, and an American Cancer Society Research Scholar Award (RSG-22-117-01-DMC) to KR.

## Supporting Information

Additional experimental and simulated data, complete strain list, and effects of analysis parameters. Raw PAmCherry-Clr4 fluorescence imaging data corresponding to the trajectories in the example *S. pombe* nucleus in the main text.

