## Supplemental Information for "Substrate recognition, not sequestration, drives the engagement of an H3K9 methyltransferase in living cells"

**Contents:**

- Supplementary Figures S1 – S10
- Supplementary Tables S1 – S5
- Supplementary Methods
- Supplementary Equation S1
- Supplementary References
- Supplementary Movie S1 Caption

### Supplementary Figures

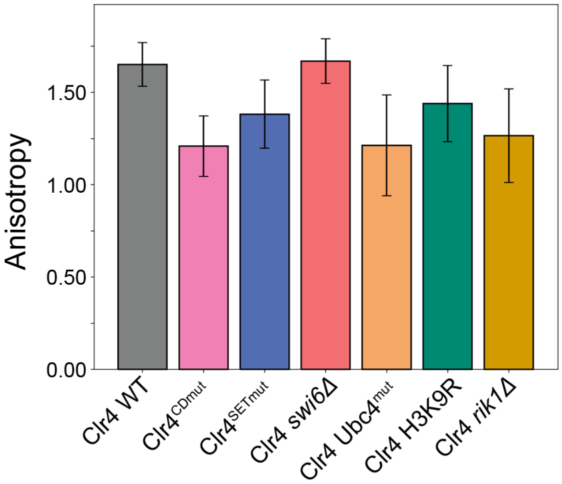


**Figure S1. Anisotropy values ($f_{180/0}$) for intermediate-range displacements (between 0.035 – 0.175 µm).** Anisotropy was assessed by comparing the fraction of angles within the range 180° ± 30° against those within 0° ± 30° across multiple length scales, i.e.,:

$$f_{180/0} = \frac{180^{\circ} \pm 30^{\circ}}{0^{\circ} \pm 30^{\circ}}$$

Error bars indicate the 95% confidence intervals from 10000 iterations of bootstrapping 100% of the data.

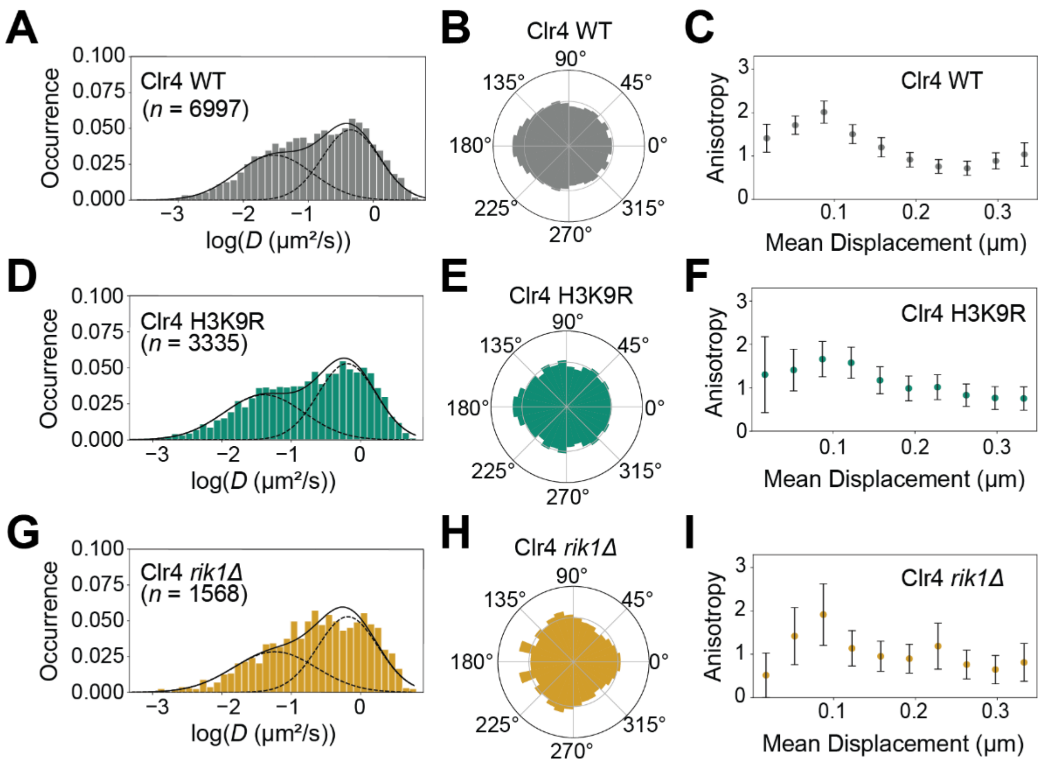


**Figure S2. PAmCherry-Clr4 single-molecule dynamics in additional mutant strains.**

**(A,D,G)** Diffusion coefficient distribution of *n* PAmCherry-Clr4 trajectories in each indicated *S. pombe* strain. Distributions are fit to a two-term Gaussian mixture model (dashed lines) to estimate the fast and slow diffusion populations (Table S2). *n* denotes the number of trajectories measured in each condition.

**(B,E,H)** Polar histograms of angles from all selected mobile single-molecule trajectories ($D_{app} \geq 0.1$ μm$^2$/s) in panels A,D,G.

**(C,F,I)** Anisotropic diffusion measurement ($f_{180/0}$) of the fast population of diffusing PAmCherry-Clr4 molecules in each strain ($D_{app} \geq 0.1$ μm$^2$/s). Error bars indicate the 95% confidence intervals from bootstrapping 100% of the data across 10000 iterations.

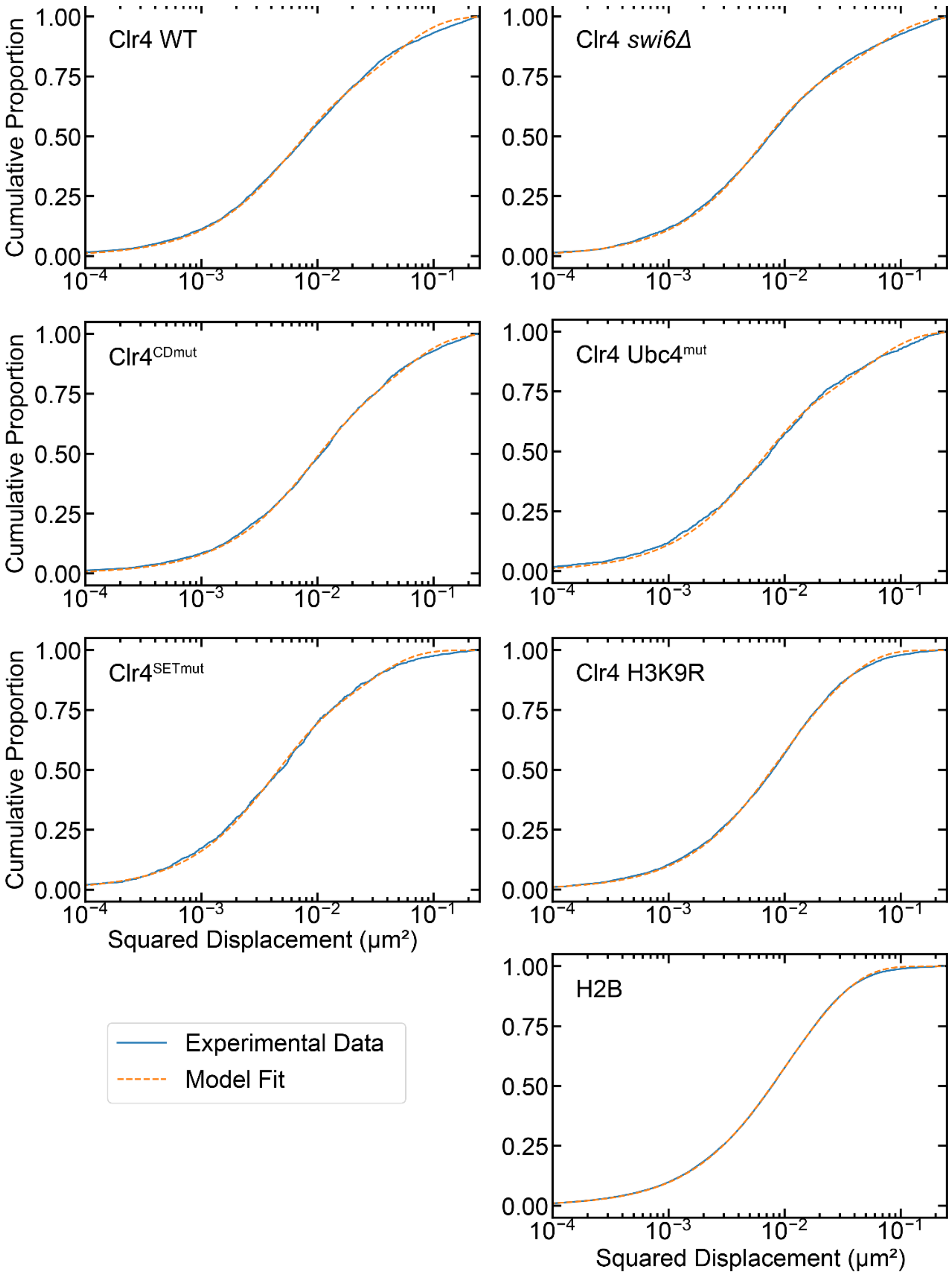


**Figure S3. Cumulative distribution functions used to determine the population weights for the time-lapse data in Figure 5.** Each distribution was fit to main text equations (3) and (4).

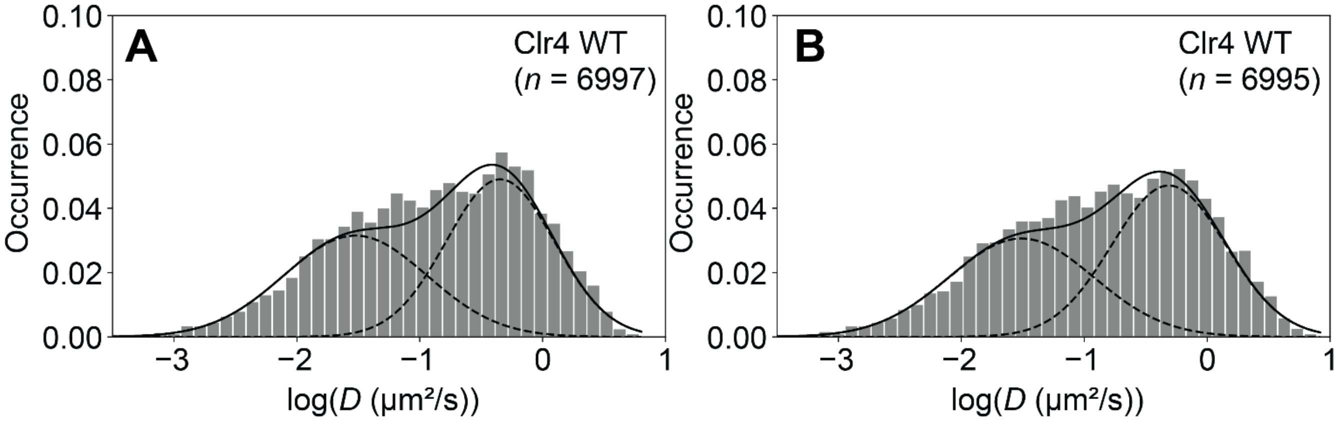


**Figure S4. Accounting for explicit motion blur correction does not change the apparent diffusion coefficient distribution of *n* PAmCherry-Clr4 trajectories analyzed.** Distributions are fit to a two-term Gaussian mixture model (dashed lines) to estimate the fast and slow diffusion populations.

**(A)** The apparent diffusion coefficient is calculated based on a blur correction that only accounts for localization uncertainty (main text Equation 1).

**(B)** The apparent diffusion coefficient is calculated based on a blur correction that accounts for localization uncertainty and the finite camera exposure time (Equation S1).

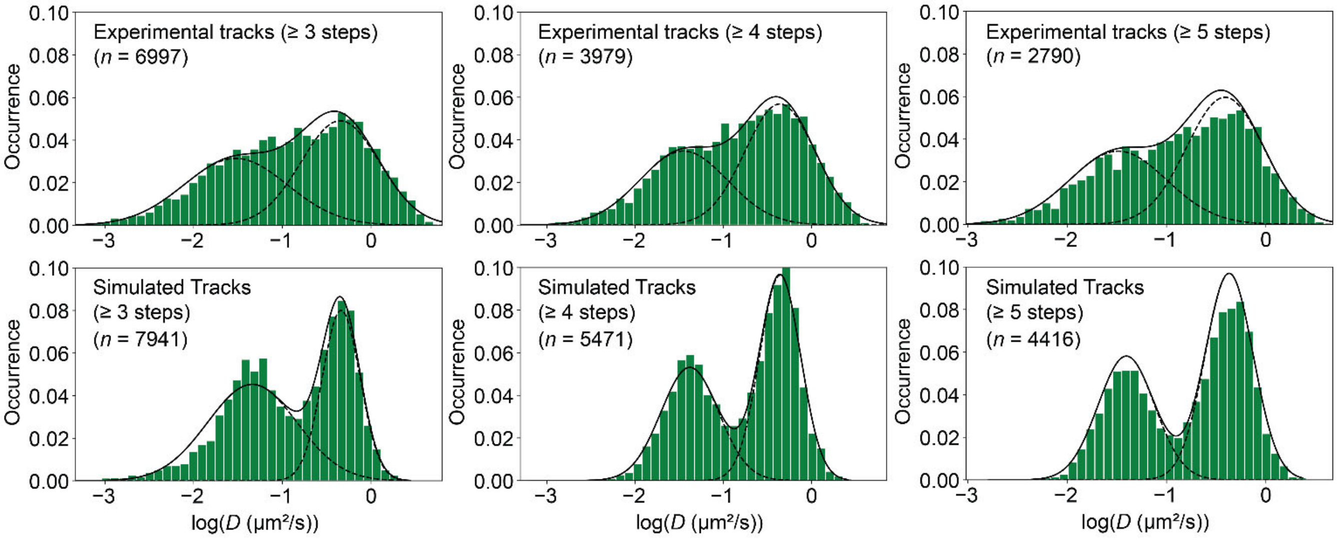


**Figure S5. Comparison of simulated and experimental data to determine the accuracy of MSD calculations.** The apparent diffusion coefficient distributions of *n* experimental or simulated PAmCherry-Clr4 trajectories are fit to a two-term Gaussian mixture model (dashed lines) to estimate the fast and slow diffusion populations. The simulated and experimental track distributions peak at similar apparent diffusion coefficients, though the peak widths are narrower for the simulated data. This analysis is largely insensitive to the minimum track length analyzed. See Table S3 for the numerical values.

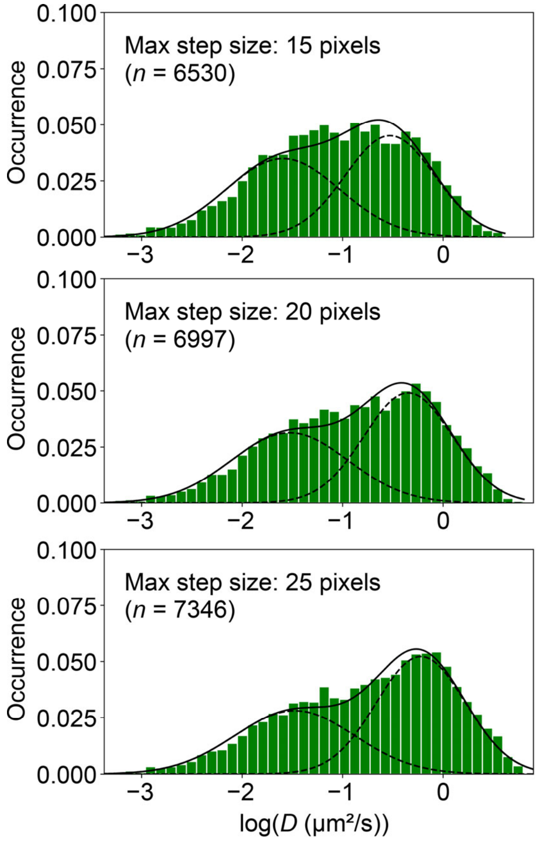


**Figure S6. Sensitivity analysis shows that the inferred fractions are robust to reasonable changes in the maximum step length (1 pixel = 49 nm).** The apparent diffusion coefficient distributions of *n* PAmCherry-Clr4 trajectories are fit to a two-term Gaussian mixture model (dashed lines) to estimate the fast and slow diffusion populations. See Table S4 for numerical values.

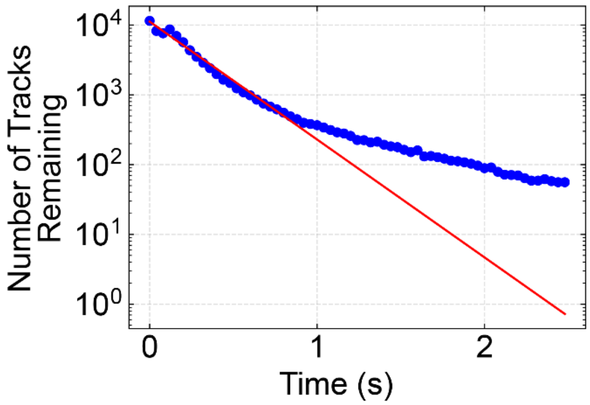


**Figure S7. Estimated probability of bleaching from fitting a single-exponential decay to fast-tracking track survival numbers.** Trajectories were simulated using the built-in function *simulate_tracks.sim_FOV* in exTrack. To accurately simulate photobleaching, we fit the measured photobleaching rate for wild-type PAmCherry-Clr4 to a single-exponential decay function. The fit is good up to 20 imaging frames (0.8 s). Blue line: experimental data. Red line: single-exponential fit.

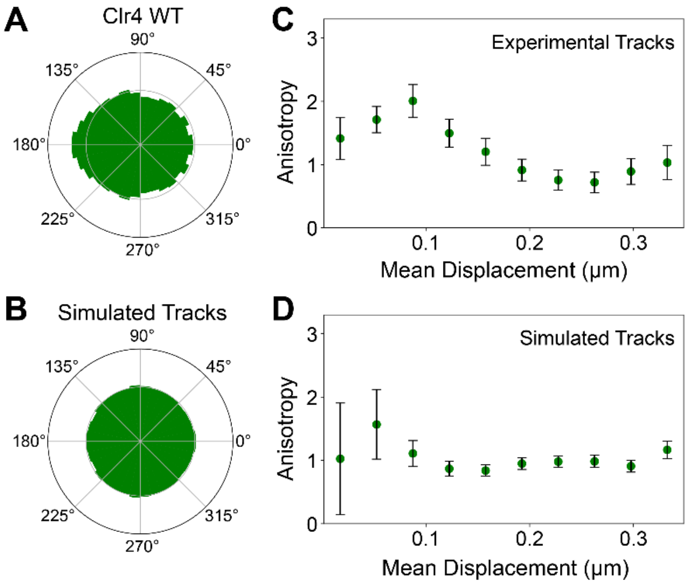


**Figure S8. Anisotropy for our experimental data and the simulated tracks.**

**(A,B)** Polar histograms of angles from all selected mobile single-molecule trajectories ($D_{app}$ ≥ 0.1 μm$^2$/s) from experimental and simulated WT PAmCherry-Clr4 tracks. The simulated tracks are almost perfectly isotropic.

**(C,D)** Anisotropic diffusion measurement ($f_{180/0}$) of the fast population of diffusing PAmCherry-Clr4 molecules in each strain ($D_{app}$ ≥ 0.1 μm$^2$/s) for experimental and simulated tracks. Error bars indicate the 95% confidence intervals from bootstrapping 100% of the data across 10000 iterations.

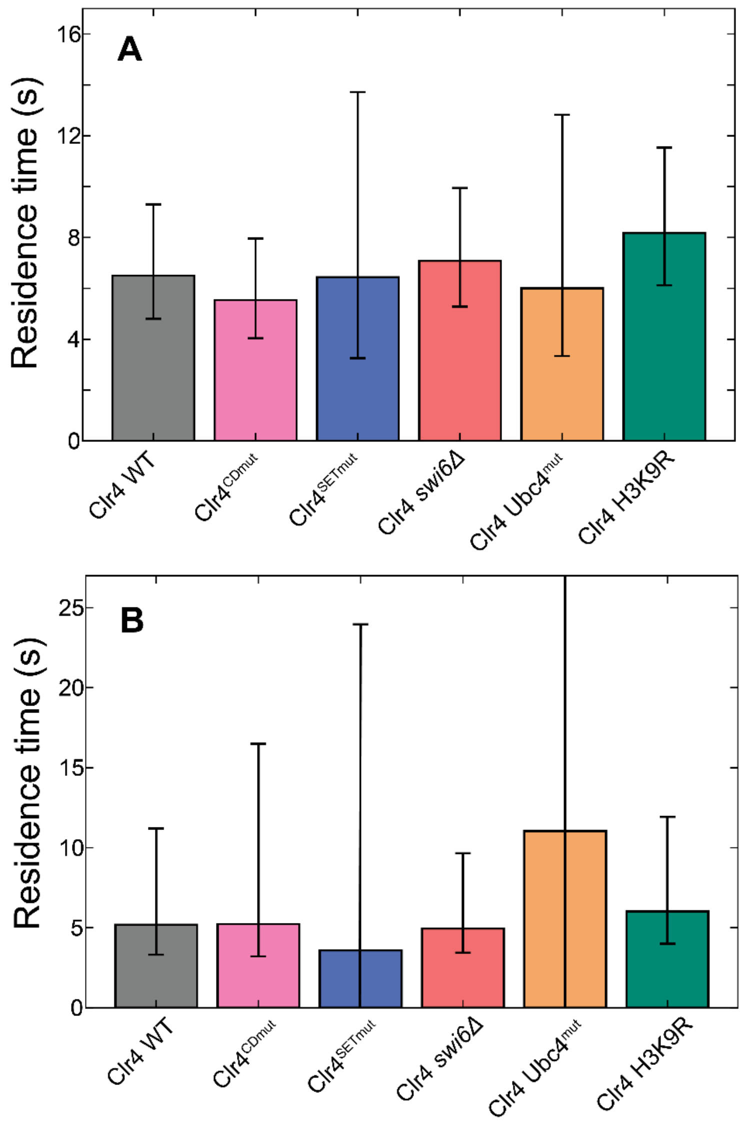


**Figure S9. PAmCherry-Clr4 residence times calculated with (A) amplitudes fixed according to the CDF and (B) amplitudes that were allowed to vary.** Each adjusted residence time is calculated from a fit to the corresponding decay curve and corrected based on the apparent decay of the nominally immobile H2B-PAmCherry. 95% confidence intervals were calculated by fitting a double-exponential decay function to 100% of the data bootstrapped 10000 times. We cannot definitively state whether residence times remain constant between methods because of the large confidence intervals in (B).

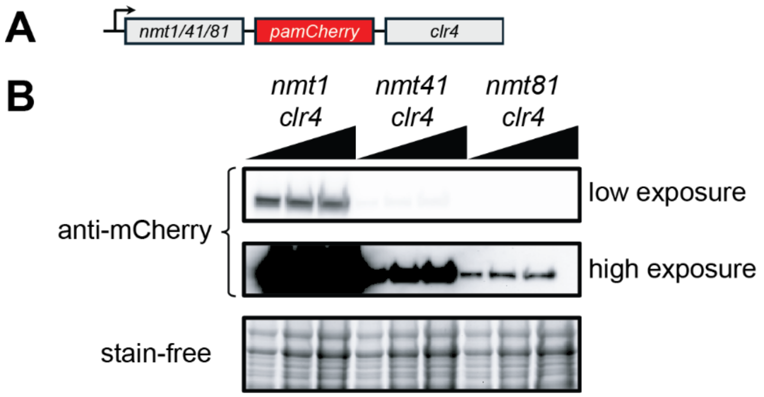


**Figure S10. PAmCherry-Clr4 expression levels.**

**(A)** PAmCherry is fused to the N-terminus of Clr4 and expressed ectopically using an inducible promoter: *nmt1, nmt41,* or *nmt81* (high, medium, or low expression levels, respectively).

**(B)** The expression levels of PAmCherry-Clr4 under *nmt1*, *nmt41*, or *nmt81* promoters were determined by western blot against an mCherry antibody. The expression levels are compared to PAmCherry-Clr4 expressed from the native Clr4 promoter.

### Supplementary Tables

#### Table S1. Strain List

| Strain Number | Genotype | Source | Related to |
|---|---|---|---|
| 683 | *h90 leu1-32 his2- ura4 DS/E ade6-M210 Kint2::ura4+* | Bao et al., 2019 (SI refs) | Fig. 2 |
| 684 | *h90 leu1-32 his2- ura4 DS/E ade6-M210 Kint2::ura4+; clr4Δ::kanMX6* | Bao et al., 2019 (SI refs) | Fig. 2 |
| 2984 | *h90 his2- ura4 DS/E ade6-M210 Kint2::ura4+ clr4Δ::kanMX6 leu1+:nmt1-PAmCherry-clr4* | This study | Fig. 2 |
| 2986 | *h90 his2- ura4 DS/E ade6-M210 Kint2::ura4+ clr4Δ::kanMX6 leu1+:nmt41-PAmCherry-clr4* | This study | Fig. 2 |
| 2988 | *h90 his2- ura4 DS/E ade6-M210 Kint2::ura4+ clr4Δ::kanMX6 leu1+:nmt81-PAmCherry-clr4* | This study | Fig. 2 |
| 2854 | *h90 ade6-M216 ura4-D18 leu1+:nmt41-PAmcherry-clr4* | This study | Figs. 3,4,5,S2 |
| 4501 | *h+ otr1R(SphI)::ade6+ ura4-D18 leu1-32 ade6-M210 clr4Δ::ura4-kanMX6 leu1+:nmt41-PAmcherry-clr4 W31G* | This study | Figs. 3,5 |
| 4500 | *h+ otr1R(SphI)::ade6+ ura4-D18 leu1-32 ade6-M210 clr4Δ::ura4-kanMX6 leu1+:nmt41-PAmcherry-clr4 H410L C412A* | This study | Figs. 3,5 |
| 4495 | *h+ otr1R(SphI)::ura4+ ura4-DS/E ade6-M210 swi6Δ::natMX6 leu1+:nmt41-PAmCherry-clr4* | This study | Figs. 4,5 |
| 4494 | *h+ otrIR(SphI)::ade6 ade6-DN/N ura4-DS/E ubc4-G48D-kanMX6 leu1+:nmt41-PAmcherry-clr4* | This study | Figs. 4,5 |
| 2251 | *h90 ade6-M216 ura4D18 leu1+:htb1-pamcherry* | Biswas et al., 2022 (ref. 16) | Fig. 5 |
| 4513 | *h+ otr1R(SphI)::ade6+ ura4-D18 ade6-M210 H3.1/H3.2/H3.3 K9R leu1+:nmt41-PAmCherry-clr4* | This study | Fig. S2 |
| 4515 | *h+ imrR(NcoI)::ura4+ ade6-216 ura4DS/E oriI rik1Δ::kanMX6 leu1+:nmt41-PAmCherry-clr4* | This study | Fig. S2 |

**Table S2. Summary of mobility measurements from single-molecule tracking measurements.** Each diffusion coefficient histogram was fit to a Gaussian mixture model to determine the apparent diffusion coefficient, *D*, and fraction, *π*, of the slow and fast states, respectively. The ranges indicate the 95% confidence intervals from bootstrapping 100% of the data across 10000 iterations.

| Strain | $D_{slow}$ (μm²/s) | $D_{fast}$ (μm²/s) | $\pi_{slow}$ (%) | $\pi_{fast}$ (%) |
|---|---|---|---|---|
| Clr4 in WT | 0.030 ± 0.002 | 0.45 ± 0.03 | 46 ± 2 | 54 ± 2 |
| $Clr4^{CDmut}$ | 0.06 ± 0.01 | 0.72 ± 0.05 | 33 ± 4 | 67 ± 3 |
| $Clr4^{SETmut}$ | 0.04 ± 0.01 | 0.57 ± 0.05 | 31 ± 4 | 69 ± 4 |
| Clr4 in *swi6Δ* | 0.027 ± 0.002 | 0.47 ± 0.03 | 44 ± 2 | 56 ± 2 |
| Clr4 in $Ubc^{mut}$ | 0.033 ± 0.009 | 0.6 ± 0.1 | 48 ± 7 | 52 ± 5 |
| Clr4 in H3K9R | 0.040 ± 0.004 | 0.64 ± 0.05 | 45 ± 3 | 55 ± 3 |
| Clr4 in *rik1Δ* | 0.06 ± 0.01 | 0.65 ± 0.09 | 43 ± 5 | 57 ± 5 |

**Table S3: Summary of mobility results in Figure S5 for analysis as a function of minimum track length.** Each diffusion coefficient histogram was fit to a Gaussian mixture model to determine the apparent diffusion coefficient, *D*, and fraction, *π*, of the slow and fast states. The ranges indicate the 95% confidence intervals from bootstrapping 100% of the data across 10000 iterations.

| **Min. track length** | $\pi_{slow}$ **(%)** | $D_{slow}$ **(μm²/s)** | $\pi_{fast}$ **(%)** | $D_{fast}$ **(μm²/s)** |
|---|---|---|---|---|
| 3 steps (experimental) | 46 ± 2 | 0.030 ± 0.002 | 54 ± 2 | 0.45 ± 0.03 |
| 3 steps (simulated) | 57 ± 2 | 0.046 ± 0.003 | 43 ± 2 | 0.46 ± 0.01 |
| 4 steps (experimental) | 43 ± 3 | 0.035 ± 0.004 | 57 ± 2 | 0.44 ± 0.03 |
| 4 steps (simulated) | 44 ± 2 | 0.042 ± 0.002 | 57 ± 2 | 0.44 ± 0.01 |
| 5 steps (experimental) | 42 ± 3 | 0.033 ± 0.004 | 58 ± 3 | 0.39 ± 0.03 |
| 5 steps (simulated) | 41 ± 2 | 0.039 ± 0.001 | 59 ± 2 | 0.43 ± 0.01 |

**Table S4. Summary of mobility results in Figure S6 for analysis after changing the maximum step size for single-molecule tracking analysis from 20 pixels (default) to 15 or 25 pixels (1 pixel = 49 nm).** Each diffusion coefficient histogram was fit to a Gaussian mixture model to determine the apparent diffusion coefficient, *D*, and fraction, *π*, of the slow and fast states, respectively. The ranges indicate the 95% confidence intervals from bootstrapping 100% of the data across 10000 iterations.

| **Max. step size (pixels)** | $\pi_{slow}$ **(%)** | $D_{slow}$ **(μm²/s)** | $\pi_{fast}$ **(%)** | $D_{fast}$ **(μm²/s)** |
|---|---|---|---|---|
| 15 | 50 ± 2 | 0.026 ± 0.002 | 50 ± 3 | 0.30 ± 0.02 |
| 20 (default) | 46 ± 2 | 0.030 ± 0.002 | 54 ± 2 | 0.45 ± 0.03 |
| 25 | 42 ± 2 | 0.033 ± 0.004 | 58 ± 2 | 0.60 ± 0.04 |

**Table S5. Summary of decay rates and residence times in main text Figure 5.** The ranges indicate the 95% confidence interval from bootstrapping 100% of the data across 10000 iterations.

| Strain | Uncorrected $k_{sb}$ ($s^{-1}$) | Corrected residence time (s) |
|---|---|---|
| Clr4 in WT | 0.34 – 0.44 | 4.8 – 9.3 |
| Clr4 in *swi6Δ* | 0.33 – 0.42 | 5.2 – 10.0 |
| Clr4 in $Ubc^{mut}$ | 0.31 – 0.53 | 3.3 – 12.9 |
| $Clr4^{SETmut}$ | 0.30 – 0.54 | 3.2 – 13.8 |
| Clr4 in H3K9R | 0.32 – 0.39 | 6.1 – 11.6 |
| $Clr4^{CDmut}$ | 0.36 – 0.47 | 4.0 – 7.9 |

### Supplementary Methods:

We used the following non-default parameters for the SMALL-LABS single-molecule tracking (Isaacoff et al. 2019):

dfrlmsz: 9
avgwin: 100
moloffwin: 200
bgsub: false
mask_fname: true
trackparams:
    minimum merit: 0.1
    integration time (ms): 40
    gamma: 0.5
    maximum step size (px): 20
    minimum track length (frames): 4
    speed estimation window halfsize: 1
    time delay between consecutive frames (ms): 0
bpthrs: 99
maxerr: .05
stdtol: 2.5
usegpu: false

### Supplementary Equation:

The equation below represents a model that fully accounts for motion blur during a single acquisition frame (Deschout et al. 2012):

$$MSD = 4D_{app}\tau + 4\sigma^2 - \frac{4}{3} D_{app} \times 0.04 \text{ sec} \quad \text{(S1)}$$

In this equation, the mean square displacement, *MSD*, is related to the apparent diffusion coefficient, $D_{app}$, the time lag, $\tau$, and the localization precision, $\sigma$.


### Supplementary Movie Caption

**Movie S1.** Raw PAmCherry-Clr4 fluorescence imaging data corresponding to the trajectories in the example *S. pombe* nucleus in main text Figure 1B,C. The cyan circle indicates the approximate nucleus position. The movie plays in real time (25 frames/sec). Scale bar: 1 μm.
